# Detection of classic and cryptic *Strongyloides* genotypes by deep amplicon sequencing: A preliminary survey of dog and human specimens collected from remote Australian communities

**DOI:** 10.1101/549535

**Authors:** Meruyert Beknazarova, Joel L. N. Barratt, Richard S. Bradbury, Meredith Lane, Harriet Whiley, Kirstin Ross

**Affiliations:** Environmental Health, Flinders University, South Australia; Division of Parasitic Diseases and Malaria, Centers for Disease Control and Prevention; Oak Ridge Associated Universities, Oak Ridge, Tennessee; Synergy America Inc.

**Keywords:** *Strongyloides*, genotypes, dogs, Australia, zoonoses, hookworms, *Ancylostoma*

## Abstract

Strongyloidiasis is caused by the human infective nematodes *Strongyloides stercoralis, Strongyloides fuelleborni* subsp. *fuelleborni* and *Strongyloides fuelleborni* subsp. *kellyi*. The zoonotic potential of *S. stercoralis* and the potential role of dogs in the maintenance of strongyloidiasis transmission has been a topic of interest and discussion for many years. In Australia, strongyloidiasis is prevalent in remote socioeconomically disadvantaged communities in the north of the continent. Being an isolated continent that has been separated from other regions for a long geological period, description of the diversity of Australian *Strongyloides* genotypes adds to our understanding of the genetic diversity within the genus. Using PCR enrichment combined with Illumina sequencing technology, we sequenced the *Strongyloides SSU* 18S rDNA hyper-variable I and hyper-variable IV regions using *Strongyloides*-specific primers, and a fragment of the mtDNA *cox*1 gene using primers that are broadly specific for *Strongyloides* sp. and hookworms. These loci were amplified from DNA extracted from Australian human and dog faeces, and one human sputum sample. Using this approach, we confirm for the first time that potentially zoonotic *S. stercoralis* genotypes are present in Australia, suggesting that dogs represent a potential reservoir of human strongyloidiasis in remote Australian communities.

**Author summary:** *Strongyloides stercoralis* is a soil-transmitted nematode that causes the disease strongyloidiasis. Due to the autoinfective nature of this parasite, it can re-infect a host causing chronic infection. If not diagnosed and treated it can be highly detrimental to human health and has a high mortality rate. Strongyloidiasis is common in remote communities in the north of Australia and has been an issue for decades. Despite various successful intervention programs to treat human strongyloidiasis, the disease remains endemic in those communities. Here for the first time we looked at the Australian dogs’ potential to infect humans and found that they carry two genetically distinct strains of *Strongyloides* spp., one of which also infects humans. This supports the hypothesis that dogs are a potential source for human strongyloidiasis. We also found that dogs in Australia might be carrying unique haplotypes. Whether these new haplotypes are also human infective is to be confirmed by further research.

## Introduction

Strongyloidiasis is caused by the human infective nematodes *Strongyloides stercoralis, Strongyloides fuelleborni* subsp. *fuelleborni* and *Strongyloides fuelleborni* subsp. *kellyi* (Grove, 1996). Worldwide, *Strongyloides* sp. are estimated to infect up to 370 million people, predominately in socioeconomically disadvantaged communities (Olsen et al., 2009, Bisoffi et al., 2013, Beknazarova et al., 2016). While *S. stercoralis* is a globally distributed nematode, *S. f. fuelleborni* has thus far only reported in Africa and Southeast Asia and *S. f. kellyi* from Papua New Guinea (PNG) (Thanchomnang et al., 2017, Pampiglione and Ricciardi, 1971, Ashford et al., 1992).

In Australia, strongyloidiasis is prevalent in remote communities located across the Northern Territory, Queensland, Western Australia, northern South Australia and northern New South Wales (Page et al., 2016). Current estimates of infection rates in some communities are up to 41% or 60% based on microscopy or serology respectively (Heydon and Green, 1931, Sampson et al., 25-26 June 2003, Miller et al., 2018, Holt et al., 2017). Despite initially successful intervention programs targeting treatment to eliminate human strongyloidiasis in remote Australian communities, the disease remains endemic (Kearns et al., 2017, Page et al., 2006). Reappearance of the infection could possibly be as a result of zoonotic transmission from dog reservoirs given that dogs and humans share a close and intimate cultural bond in rural and remote Indigenous communities of Australia (Constable et al., 2010).

The zoonotic potential of *S. stercoralis* infected dogs and their potential role in the maintenance of strongyloidiasis transmission has been a topic of interest and discussion for many years (Beknazarova et al., 2017b, Goncalves et al., 2007, Takano et al., 2009). Molecular investigation of human and dog derived *S. stercoralis* is useful in understanding the nature of cross infection. There are two regions of the *S. stercoralis* nuclear and mitochondrial genome that are considered to be conserved within the *Strongyloides* genus and can be used as markers for molecular typing of *S. stercoralis*. Hyper-variable regions (HVR) I and IV of the small Subunit (*SSU*) 18s ribosomal DNA and the cytochrome c-oxidase subunit 1 (*cox*1) gene of the mitochondrial DNA (mtDNA) have been widely used to study relationships between *S. stercoralis* from different hosts and different geographic locations (Hasegawa et al., 2009, Hasegawa et al., 2010, Jaleta et al., 2017, Nagayasu et al., 2017, Basso et al., 2018). Based on genetic analysis of these loci, it has been recently found that there are two genetically different *S. stercoralis* strains, one is dog and human infective, and the other is dog specific. These data were collected from dogs and humans in Cambodia and Japan (Jaleta et al., 2017, Nagayasu et al., 2017).

Being an isolated continent that has been separated from other regions for a long geological period, Australia could represent an interesting addition to our understanding of the genetic diversity within *S. stercoralis* and the *Strongyloides* genus more generally. Indigenous Australians have inhabited the continent for at least 40,000 years and dogs (in the form of dingoes) were likely introduced up to 12,000 years ago (Clutton-Brock, 1995). Given this long period of relative isolation, it might be expected that Australia could harbor unique endemic genotypes or unique sub-species of *S. stercoralis* that have evolved within dog and human populations over this period.

Using PCR enrichment combined with Illumina sequencing technology, we sequenced the *Strongyloides SSU* 18S rDNA hyper-variable I and hyper-variable IV regions using *Strongyloides*-specific primers, and a fragment of the mtDNA *cox*1 gene using primers that are broadly specific for *Strongyloides* sp. and hookworms. This approach was applied to DNA extracted from human and dog faeces, and one human sputum sample. The main focus of this study was to genotype Australian human and dog *S. stercoralis* strains to see whether dogs carry human *S. stercoralis* strains and/or vice versa. To our knowledge this is the first time human and dog *S. stercoralis* have been studied in Australia on a molecular level.

## Methods

### Study area and faeces collection

Dog faecal samples were collected from communities in the Northern Territory, Australia. Dog faeces were collected from the environment (i.e., the ground) in the selected communities and stored in the DESS (dimethyl sulfoxide, disodium EDTA, and saturated NaCL) solution to preserve the DNA (Beknazarova et al., 2017a). For those dog faeces that were collected from the private land, the consent forms were received to collect the samples. The preserved faecal samples were express posted to the Environmental Health laboratory, Flinders University, South Australia, for further analysis.

Human faecal and sputum samples were provided by our colleagues at the Royal Darwin Hospital, NT, AusDiagnostics Pty Ltd, NSW, and Townsville Hospital, Queensland. While the personally-identifying information of the patients was de-linked from our analyses, their infections are known to have been locally acquired. Ethics approval from the Social and Behavioural Research Ethics Committee (SBREC) No 6852 dated 1^st^ June 2015 was obtained for collecting dog faeces from the remote communities. Human ethical approval from the Southern Adelaide Clinical Human Research Ethics Committee (SAC HREC) No 309.17 dated 24^th^ January 2018 was obtained for comparing *S. stercoralis* DNA extracted from human and dog tissues. CDC investigators were not engaged with sample collection and their participation did not include engagement with human or animal subjects.

### DNA extraction

Prior to DNA extraction, faecal samples containing DESS were centrifuged for three minutes a t 3000 x g rpm using an Orbital 400 Clements (Phoenix, Lidcombe, Australia). The supernatant consisting of the preservative solution was removed. The remaining faecal sample was washed with sterile saline solution. DNA was extracted using the Power Soil DNA isolation kit (QIAGEN, Hilden, Germany) following the manufacturer’s instructions with slight modifications that included incubating samples at 56°C overnight after the cell lysis step, followed by vortexing of samples for three minutes. Approximately 250 milligrams of the pellet was placed into a PowerBead tube containing lysis buffer (included in the Power Soil DNA extraction kit). The remainder of the extraction process was performed according to the manufacturer’s instructions. Extracted DNA was stored at −20°C prior to real-time PCR (qPCR) analysis.

### Real-time PCR

The samples (273 dog and 4 human DNA samples) were first screened for *Strongyloides* spp. using qPCR in the Environmental Health laboratory at Flinders University, SA, Australia. The real-time PCR assay was adopted from Verweij *et al*. (2009) using *S. stercoralis* - specific primers targeting a 101 base pair region of 18S rRNA. The 20 µL reaction contained 10 µL Supermix (SSoAdvanced, Universal Probes Supermix, Foster City, CA, USA), 1 µL primers and probe mixture (Stro18S-1530F, Stro18S-1630R and Stro18S-1586T FAM) (Bio-Rad Laboratories, CA, USA), 4 µL deionised H2O, and 5 µL DNA template. All qPCR reactions were performed in triplicate on the Corbett Rotor-Gene 6000 machine (QIAGEN, Hilden, Germany). *S. stercoralis* primers and probes and qPCR conditions used are shown in Table 1. A sample was considered positive when the Ct value was lower than the mean negative Ct minus 2.6 standard deviations of a mean negative control Ct. Positive samples were amplified in every qPCR reaction.

### Conventional PCR for amplification of *SSU* HVR-I and HVR-IV, and *cox*1 sequences

Extracted DNA from samples that were qPCR positive for *Strongyloides* spp. (47 dog and four human DNA samples) was shipped on dry ice to the Centers for Disease Control and Prevention (CDC), Georgia, USA for conventional PCR, sequencing and bioinformatics analysis. Hyper-variable regions (HVR) I and IV in the small Subunit (*SSU*) 18S ribosomal DNA and a fragment of the mitochondrial cytochrome c-oxidase subunit 1 (*cox*1) gene were amplified using conventional PCR and then sequenced using Illumina technology. All PCR reactions were performed on a GeneAmp PCR System 9700 Thermo Cycler, version 3.12 (Applied Biosystems, USA). *S. stercoralis* primers and PCR conditions used for qPCR and conventional PCR are shown in Table 1. For the *cox*1 gene, PCR reactions were performed in a total volume of 50 µL containing 10 µL NEB 5X Q5® Buffer (New England BioLabs, USA), 10 µL NEB 5X Q5® High GC Enhancer (New England BioLabs, USA), 4 µL NEB Deoxynucleotide Solution Mix (10 mM each nt) (New England BioLabs, USA), 1 µL Q5® High-Fidelity DNA Polymerase (New England BioLabs, USA), 2.5 µL forward primer (SSP_COX1_F), 2.5 µL reverse primer (SSP_COX1_R), 18 µL deionised H2O, and 2 µL DNA template. For the HVR-I and HVR-IV regions, PCR reactions were performed in a 25 µL reaction containing 12.5 µL of NEBNext® Q5® Hot Start HiFI PCR Mastermix, MO543L (New England BioLabs, USA), 1.5 µL forward primer (NEW_HVR_I_F or NEW_HVR_IV_F), 1.5 µL reverse primer (NEW_HVR_I_R or NEW_HVR_IV_R), 7.5 µL deionised H2O, and 2 µL DNA template.

The amplified PCR products were separated by 1.5% agarose gel electrophoresis and stained with ethidium bromide. The stained DNA bands were visualised by UV illumination using a Ugenious 3 (SYNGENE, Japan). For quality control, each PCR run included a positive control containing *Strongyloides* genomic DNA as template, a non-template control containing autoclaved sterile water instead of template, and a negative control containing DNA extracted from a parasite-free specimen. Amplicons for each of the three markers were also generated for *Strongyloides ratti* as an additional control for the sequencing and *in silico* analysis steps.

**Table 1:**
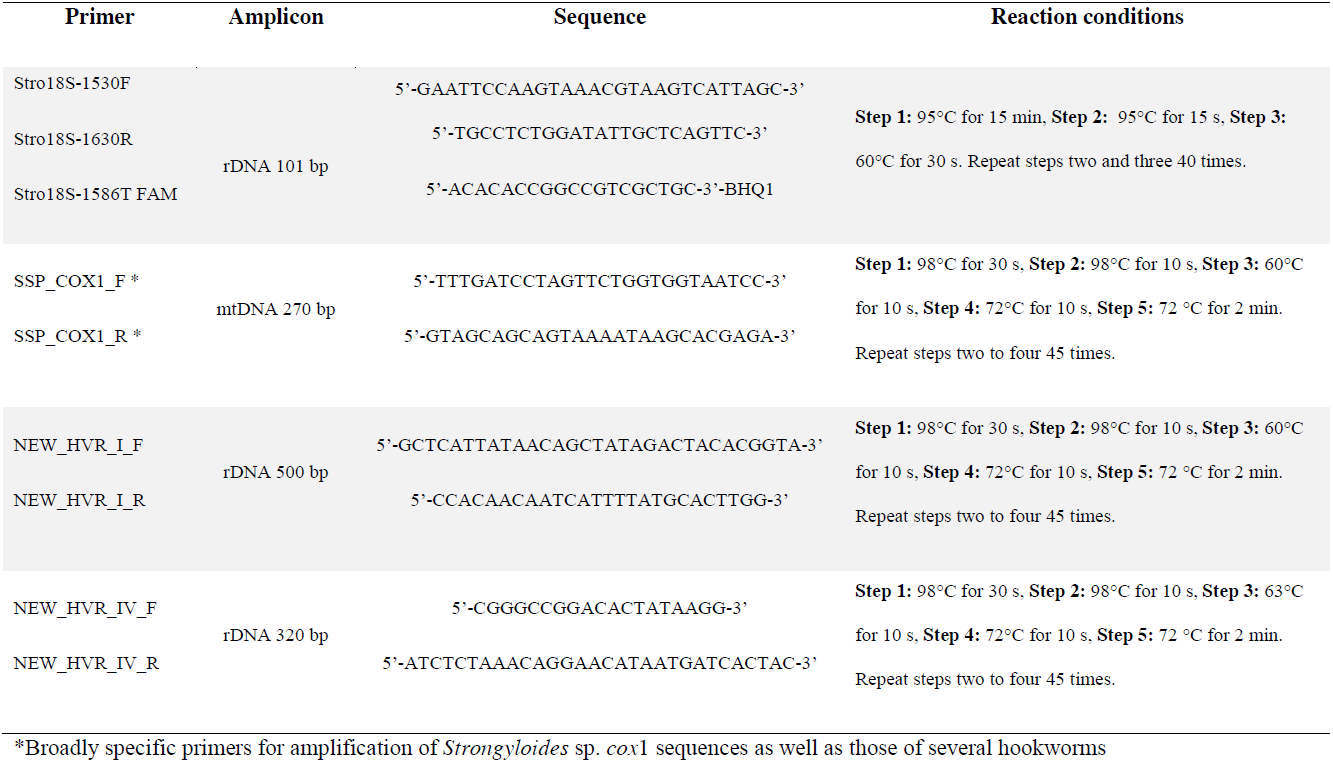
Primers and probes and PCR conditions.

### Next Generation Sequencing (NGS)

Ten microliters of PCR amplicon was purified and normalized for concentration prior to library preparation using SequalPrep Normalization Plate Kit (Thermo Fisher Scientific, USA). DNA libraries were prepared using the NEBNext Ultra DNA Library Prep Kit for Illumina (New England BioLabs, USA), and NEBNext Multiplex Oligos for Illumina Index kit (New England BioLabs, USA). The sequencing reactions were prepared using the MiSeq reagent Nano Kit v2 (PE250bp), and performed on the Illumina MiSeq platform (Illumina).

### In silico analysis

The Illumina reads were analyzed using Geneious (www.geneious.com) by means of a workflow that performed read quality control, assembly of contigs and genotype assignment. As part of this workflow, quality trimming to a minimum phred score of 20 and removal of adapter sequence was performed using BBDuk (v 37.64). Reads less than 50 base pairs (bp) in length were discarded. Paired reads were then merged using BBMerge (v 37.64) and all reads (merged and unmerged) were mapped to a reference sequence. For *cox*1 amplicons, reads were mapped to a *S. stercoralis* sequence with the GenBank (GB) accession number LC050212.1. For *SSU* HVR-I and *SSU* HVR-IV, reads were mapped to a *S. stercoralis* with the accession AF279916.2. Prior to mapping, each reference sequence was trimmed to the length of the amplicon, excluding the primer sequences. Mapping was performed using the Geneious mapper under the Medium sensitivity / Fast default settings. Reads that successfully mapped (and their corresponding mates) were retained for *de novo* assembly. These reads were assembled using the Geneious *de novo* assembler with the following customized parameters; minimum overlap: 50 bp, minimum overlap identity: 100%, maximum number of mismatches per read: 0%, and the maximum number of ambiguities: 1. Contigs were split if coverage fell below 50 bp and sub-variants with coverage less than 50 bp were not considered. Contigs were trimmed to the primer sequence using the ‘Trim Ends’ function in Geneious. Contigs were validated by taking the trimmed and merged reads and mapping them back to the contigs generated using the Geneious mapper, and the following custom parameters: minimum mapping quality: 30 bp, minimum overlap: 150 bp, minimum overlap identity: 100%, maximum number of mismatches: 0%, and the maximum number of ambiguities: 1. Contigs were split or discarded if the coverage fell below 300 to 500 bp depending on the depth obtained for a particular specimen (500 bp for specimens that obtained >10k reads, 300 for specimens with <10k reads). Finally, genotypes were assigned by performing a local BLASTN search (within Geneious) against a database constructed from all unique *Strongyloides* sp. 18S and *cox*1 sequences available in GenBank and the DNA Databank of Japan. A set of homologous sequences from several other roundworms (parasitic and free-living) were also included in this database. Sequences were only added to the BLASTN database if they overlapped our 217 bp *cox*1 amplicon by more than 95%. This included several previously published *cox*1 haplotypes that had matching *SSU* HVR­I and HVR-IV genotypes assigned (Jaleta et al., 2017).

Any specimens for which *Strongyloides ratti* sequences were detected as part of this workflow were considered to be at risk of contamination from our positive control and potentially from other specimens included in the study. Any possibly contaminated specimens were excluded from further analysis.

### Construction of a *cox*1 cluster dendrogram

Sequences of *cox*1 were exported from Geneious as a fasta file and were aligned using the ‘msa’ package in R. The ‘dist.alignment’ function from the ‘seqinr’ package was used to compute a pairwise identity matrix, considering gaps in the identity measure. The resulting matrix was clustered using the agglomerative nested clustering approach performed with the ‘agnes’ R package, using “euclidean” distances and the “average” clustering method. A cluster dendrogram was generated using the ‘ggtree’ R package. Vector images (i.e. graphics) used for annotation of the dendrogram were either generated in house at CDC or obtained from PhyloPic (http://phylopic.org).

## Results

### Real-time PCR, conventional PCR and sequencing

We screened 273 dog and four human samples using real-time PCR. Forty seven (47) dog and four human samples that were positive by qPCR were selected for conventional PCR with further sequencing of their *cox*1, *SSU* HVR-I and HVR-IV regions. In some cases, useable sequences were not generated due to poor amplification of the PCR product. Sequence data was obtained for 24 specimens including four human specimens and twenty dog specimens. The complete genotype of all specimens analyzed in this study is shown in Figure 1 and summarized in Table 2.

**Figure 1.**
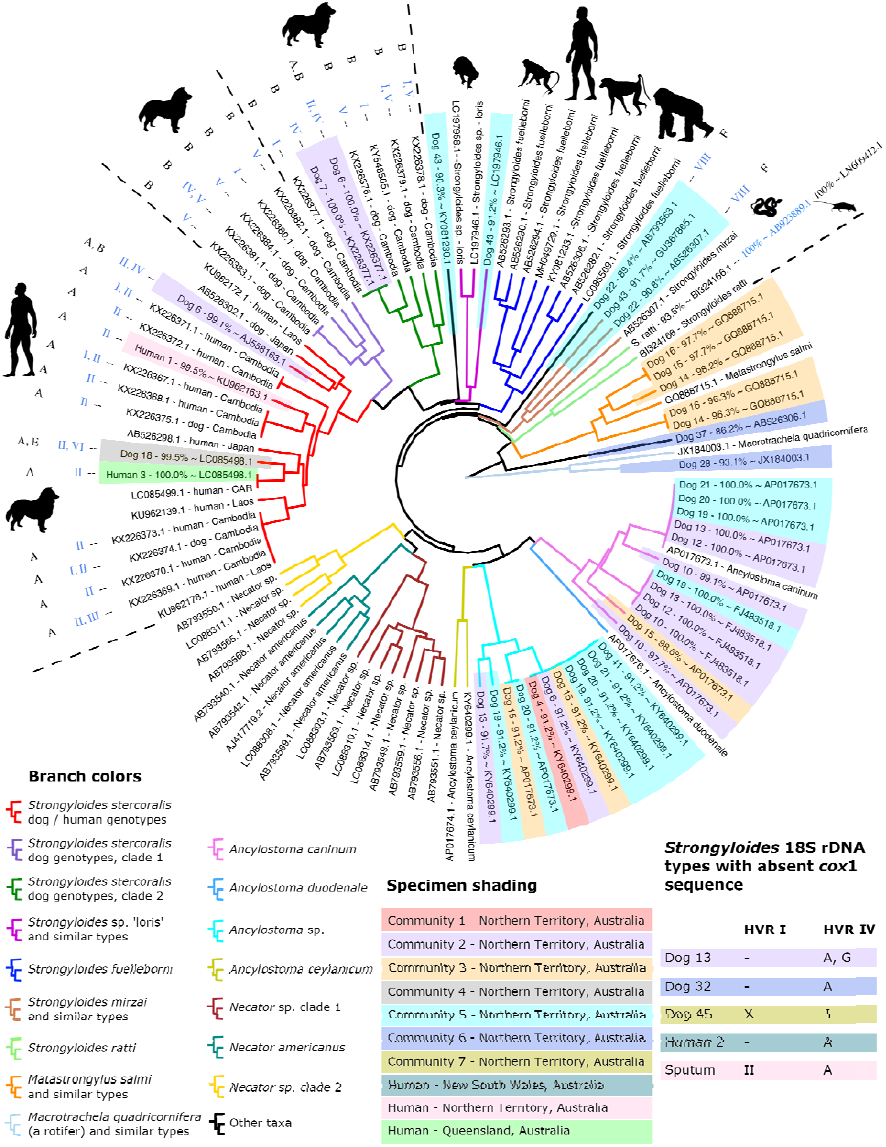
Dendrogram of clustered *cox*1 amplicons from Australian dog and human specimens. This dendrogram includes *cox*1 sequences generated in this study and a selection of previously published *cox*1 sequences that overlap with our 217 base pair *cox*1 amplicon by 100%. Specimens analyzed as part of this study are shaded according to their site of collection. Branches are color coded according to their identity; either a species assignment, a proposed genus assignment, or their *S. stercoralis* genotype. When available, *Strongyloides* sp. *cox*1 sequences are annotated with their associated *SSU* genotypes, with their HVR-I type shown in blue and their HVR-IV type shown in black. Specimens for which a *cox*1 sequence was not obtained are shown in a table embedded in the figure (bottom right), which includes two specimens possessing unique 18S genotypes; dog 13 (HVR-IV, type G) and dog 45 (HVR-I, type X). A dash (-) shown in this table indicates failed amplification and/or sequencing of that marker. ‘Sputum’ refers to the sole sputum sample from a human patient (human 4) included in this study. Sequences published in previous studies that are not from *S. stercoralis* are labelled with their GenBank and/or DNA Data Bank of Japan accession numbers followed by their species name. *Strongyloides stercoralis* sequences from previous studies are labelled with their accession number, host species, and country of origin. Note that ‘CAR’ means Central African Republic. Names of specimens collected as part of this study begin with a host name and a unique number assigned in this study, followed by a percentage similarity to (∼) a near BLASTN hit identifiable by its accession number.

**Table 2:**
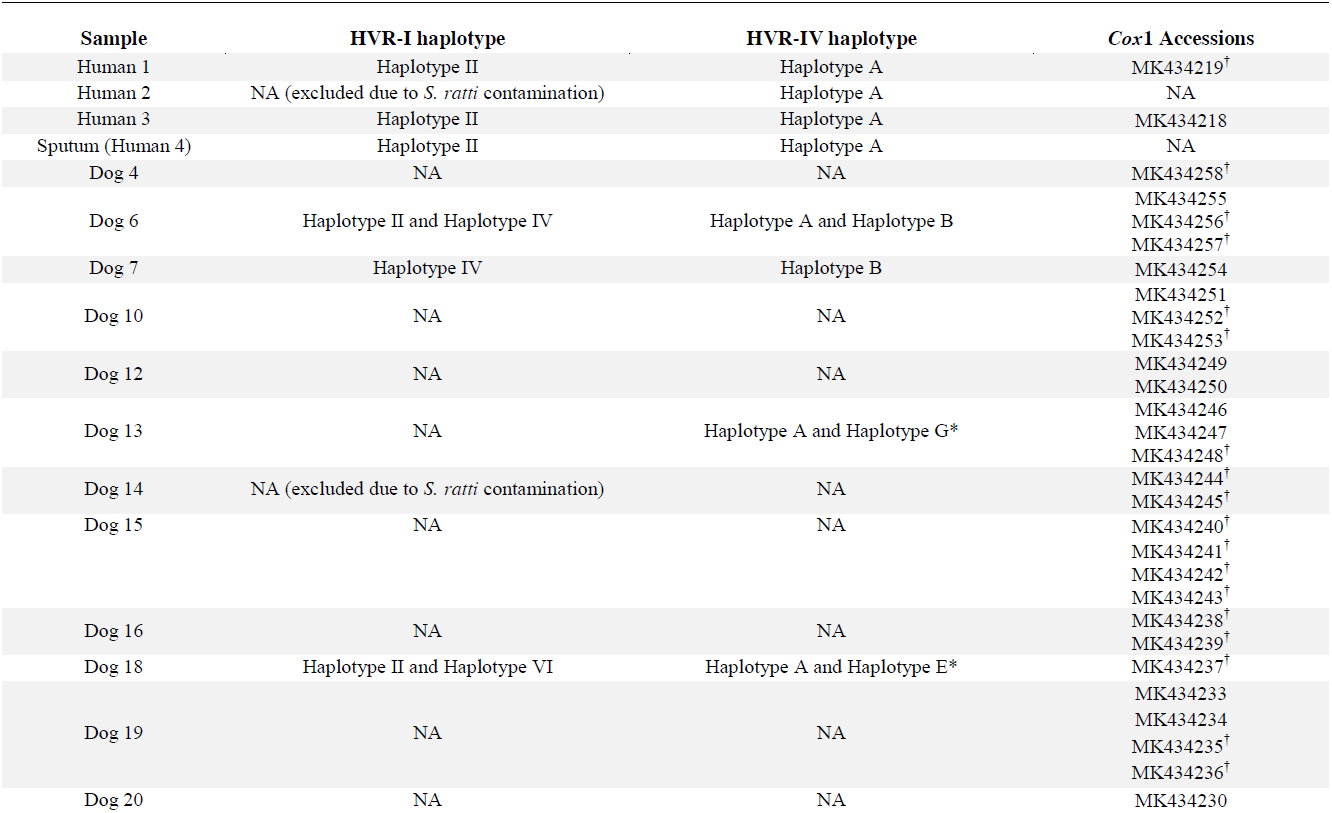
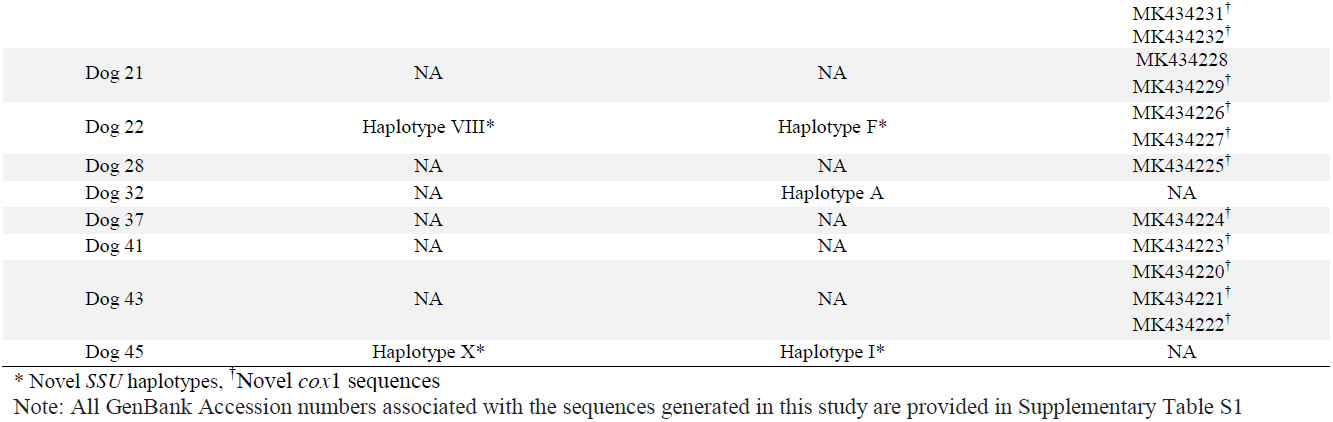
Human and dog samples analyzed in this study and their complete genotype.

| Sample | HVR-I haplotype | HVR-IV haplotype | Cox1 Accessions |
| --- | --- | --- | --- |
| Human 1 | Haplotype II | Haplotype A | MK434219<sup>†</sup> |
| Human 2 | NA (excluded due to *S. ratti* contamination) | Haplotype A | NA |
| Human 3 | Haplotype II | Haplotype A | MK434218 |
| Sputum (Human 4) | Haplotype II | Haplotype A | NA |
| Dog 4 | NA | NA | MK434258<sup>†</sup> |
| Dog 6 | Haplotype II and Haplotype IV | Haplotype A and Haplotype B | MK434255<br>MK434256<sup>†</sup><br>MK434257<sup>†</sup> |
| Dog 7 | Haplotype IV | Haplotype B | MK434254 |
| Dog 10 | NA | NA | MK434251<br>MK434252<sup>†</sup><br>MK434253<sup>†</sup> |
| Dog 12 | NA | NA | MK434249<br>MK434250 |
| Dog 13 | NA | Haplotype A and Haplotype G* | MK434246<br>MK434247<br>MK434248<sup>†</sup> |
| Dog 14 | NA (excluded due to *S. ratti* contamination) | NA | MK434244<sup>†</sup><br>MK434245<sup>†</sup> |
| Dog 15 | NA | NA | MK434240<sup>†</sup><br>MK434241<sup>†</sup><br>MK434242<sup>†</sup><br>MK434243<sup>†</sup> |
| Dog 16 | NA | NA | MK434238<sup>†</sup><br>MK434239<sup>†</sup> |
| Dog 18 | Haplotype II and Haplotype VI | Haplotype A and Haplotype E* | MK434237<sup>†</sup> |
| Dog 19 | NA | NA | MK434233<br>MK434234<br>MK434235<sup>†</sup><br>MK434236<sup>†</sup> |
| Dog 20 | NA | NA | MK434230 |
|  |  |  | MK434231<sup>†</sup> |
|  |  |  | MK434232<sup>†</sup> |
| Dog 21 | NA | NA | MK434228 |
|  |  |  | MK434229<sup>†</sup> |
| Dog 22 | Haplotype VIII* | Haplotype F* | MK434226<sup>†</sup> |
|  |  |  | MK434227<sup>†</sup> |
| Dog 28 | NA | NA | MK434225<sup>†</sup> |
| Dog 32 | NA | Haplotype A | NA |
| Dog 37 | NA | NA | MK434224<sup>†</sup> |
| Dog 41 | NA | NA | MK434223<sup>†</sup> |
| Dog 43 | NA | NA | MK434220<sup>†</sup> |
|  |  |  | MK434221<sup>†</sup> |
|  |  |  | MK434222<sup>†</sup> |
| Dog 45 | Haplotype X* | Haplotype I* | NA |

\* Novel *SSU* haplotypes, <sup>†</sup> Novel *cox1* sequences
Note: All GenBank Accession numbers associated with the sequences generated in this study are provided in Supplementary Table S1

### *SSU* HVR-I haplotypes detected among *S. stercoralis* from Australian dogs and humans

Jaleta *et al*. (2017) previously sequenced the *SSU* HVR-I region of *S. stercoralis* worms from Cambodian dog and human specimens and identified five different haplotypes (HVR-I haplotypes I-V) (Jaleta et al., 2017). A recent study of European dogs identified a new haplotype from the HVR-I region (haplotype VI) (Basso et al., 2018). In our Australian samples we found haplotype II in both human and dog samples and haplotype IV in dog samples only, which is consistent with the findings from Jaleta *et al*. (2017). We also identified haplotype VI in a single Australian dog. Following the genotype nomenclature developed by Jaleta *et al*. (2017) and Basso *et al*. (2018), we discovered two new HVR-I haplotypes; haplotypes VIII and X (Figure 2, Table 1), in addition to the six haplotypes previously described (Jaleta et al., 2017, Basso et al., 2018), Due to the existence of noteworthy similarities (> 99% in all cases) between sequences of *S. stercoralis*, *Strongyloides procyonis*, a sequence assigned to *Strongyloides* sp. Okayama (GB: LC038066.1), and our novel dog sequences, we expanded the Jaletta *et al*. typing scheme to include these sequences. This involved inclusion of haplotypes that could not be confidently assigned to *S. stercoralis* given the information on hand, yet are highly similar to known *S. stercoralis* 18S haplotypes. This adjustment was also required because a sequence attributed to *S. procyonis* (GB: AB272234.1) possesses HVR-I haplotype IV, which is identical to an *S. stercoralis* genotype assigned to a Cambodian dog (GB: KU724124.1). For details, refer to Figure 2 and Table 3.

**Figure 2.**
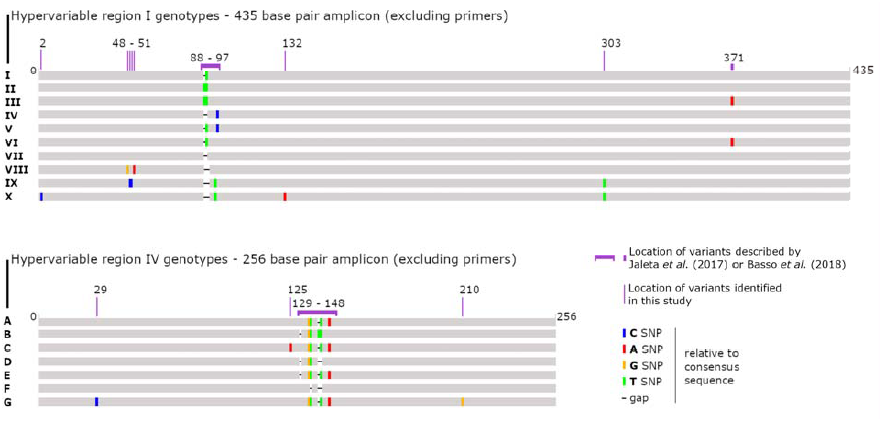
Schematic detailing the proposed modifications to the previously described ed *S. stercoralis* genotyping scheme. A graphical representation of the novel *Strongyloides* genotypes discovered in this stu udy compared to haplotypes identified in previous reports. The location of sites that wwere genotypically informative based on the original genotyping method described by Jaletta *et al*. (2017) and Basso *et al*. (2018) are indicated, as well as new SNP/indel sites that have be een incorporated into the typing scheme based on the results of this study (Jaleta et al., 2017, Ba asso et al., 2018). For hypervariable region I, we introduce two novel types (VIII and X), and assissignnew haplotype names to published sequences that had not been previously considered in th this typing scheme (VII and IX). For hypervariable region IV, we introduce three novel types (E,E, F, G and I), and assign new haplotype names to published sequences that had not been previou ously considered in this typing scheme (C, D and H) (see Tables 3 and 4 for detai ils).

**Table 3.** HVR I genotypes assigned to *Strongyloides* sp. based on current data.

| Haplotypes | GenBank Accession/s | Notes |
| --- | --- | --- |
| I | AB923888.1, AF279916.2, AJ417023.1,<br>MK468654 | Found in dogs and humans. |
| II | KF926659.1, MK468655, MK468656,<br>MK468657, MK468658 | Found in dogs and humans. |
| III | AB453315.1 | Fund in dogs and humans. |
| IV | AB272234.1, KU724124.1, MK468663 | This haplotype has been detected in dogs though not in humans. It has also been described in a badger where it was assigned to *S. procyonis*. |
| V | Jaleta *et al.* (2017) | Described by Jaleta *et al.* (2017) but the sequence of this type was not available in GenBank prior to this study. This is potentially a dog-specific haplotype. |
| VI | AB453316.1, AB453314.1, MH932098.1,<br>MH932099.1, MH932100.1, MK468660 | Assigned only to *S. stercoralis*. Identified in dog 18 from this study. Described predominantly in dogs but also in a chimpanzee (AB453314.1). |
| VII | AB205054.1 | A *S. procyonis* sequence greater than 99% similar to *S. stercoralis* Haplotype I and Haplotype IV. |
| VIII | MK468661 | Novel *Strongyloides* sp. sequence from Dog 22 most similar to Haplotype VII. |
| IX | LC038066.1 | *Strongyloides* sp. Okayama isolated from a Japanese striped snake. This sequence is greater than 99% identical to sequences of *S. stercoralis* and *S. procyonis*. Similar to Haplotype X (see below) identified in an Australian dog. |
| X | MK468662 | Sequence identified in dog 45 in present study. Most similar to *Strongyloides* sp. Okayama (Haplotype IX). |

### *SSU* HVR-IV haplotypes detected among *S. stercoralis* from Australian dogs and humans

In the Australian samples we identified HVR-IV haplotype A in both humans and dogs, and haplotype B only in dogs, as previously observed (Jaleta et al., 2017). Supporting the findings of Jaleta *et al*. (2017), our results also showed that haplotype II of HVR-I is found in combination with haplotype A of HVR-IV, and haplotype IV of the HVR-I region is only found in combination with haplotype B of the HVR-IV region (Jaleta et al., 2017). We observed that a unique sequence attributed to *S. stercoralis* had been submitted to GenBank in 1993, and this was assigned to haplotype C (GB: M84229.1). Given the strong similarity between HVR-IV sequences of *S. procyonis* and *S. stercoralis* and the fact that HRV-I haplotye IV (Jaleta *et al*. 2017) is also found in *S. procyonis* SSU sequences, we assigned the HVR-IV sequence from *S. procyonis SSU* DNA to haplotype D (GB: AB272234.1 and AB205054.1). A HVR-IV genotype 99% similar to the *Strongyloides* sp. Okayama (GB: LC038066.1) was detected in dog 45. Therefore, the HVR-IV sequence of *Strongyloides* sp. Okayama was assigned to haplotype H, and the sequence from dog 45 was assigned to Haplotype I. Consequently, four new haplotypes detected in four Australian dog samples were assigned to HVR-IV haplotypes E, F, G and I (Figure 1, Table 4).

**Table 4.** HVR IV genotypes assigned to Strongyloides sp. based on current data.

| Haplotypes | GenBank Accession/s | Notes |
| --- | --- | --- |
| A | KY081221.1, KU724128.1, KU724125.1, LC085483.1, LC085482.1, LC085481.1, KU962182.1, KU962181.1, KU962180.1, KU962179.1, AB923888.1, KF926662.1, KF926661.1, AF279916.2, MH932097.1, MH932097.1, MH932095.1, KY081223.1, AB526826.1, AB453316.1, AB453315.1, AB453314.1, MK468664 - MK468671 | Identified in dogs, humans and chimpanzees. A *S. stercoralis*- specific haplotype |
| B | KU724129.1 | Identified only in dogs. Consistently assigned to *S. stercoralis* |
| C | M84229.1 | This sequence was published in GenBank in 1993 and assigned to *S. stercoralis*. It has not appeared in the literature since based on our knowledge, though we assigned to genotype C for its historic value. It shares one SNP difference to type A (Figure 2). |
| D | AB272234.1, AB205054.1 | Includes two sequences assigned to *S. procyonis*. This type is 99% similar to Haplotype B. |
| E | MK468674 | Novel genotype identified in dog 18. Most similar to Haplotype A. |
| F | MK468675 | Novel genotype identified in dog 22. Most similar to Haplotype D. |
| G | MK468676 | Novel genotype identified in dog 13. Most similar to Haplotype A. |
| H | LC038066.1 | *Strongyloides* sp. Okayama isolated from a Japanese striped snake. |
| I | MK468677 | Novel genotype identified in dog 45. Most similar to Haplotype H. |

### Clustering of *Strongyloides stercoralis* based on *cox*1 sequences

A 217 base pair fragment of *cox*1 was sequenced from 20 Australian specimens including those from 18 dogs and two humans, plus the *S. ratti* control (21 *cox*1 sequences in total). Multiple *cox*1 types were obtained from a single specimen in many cases, revealing infections caused by multiple helminth species and multiple *S. stercoralis* genotypes in a single host (Figure 1). Dendrogram construction by agglomerative nested clustering revealed three distinct *S. stercoralis* clades, including one occupied predominantly by worms possessing the II/A *SSU* genotype, which constituted sequences obtained from dogs and humans. Four *cox*1 sequences obtained in this study (one from each of human 1, human 3, dog 6, and dog 18), were assigned to the dog and human-infecting *S. stercoralis* clade. A *S. stercoralis* clade occupied mostly by specimens possessing the I/B and V/B *SSU* genotypes was also apparent (one specimen possessed the IV/B genotype), representing dog infections only (dog clade 1). None of the Australian specimens were assigned to this clade (Figure 1). A final *S. stercoralis* clade containing *cox*1 sequences obtained from only dogs (dog clade 2) was also dominated by specimens possessing the I/B and V/B genotypes, though two specimens were also assigned the IV/B. A single *cox*1 sequence from each of dogs 6 and 7 was assigned to this clade (Figure 1).

### Cryptic *cox*1 sequences potentially from *Strongyloides* sp. helminths that could not be assigned to a genus or species

Sequences were obtained from two dog fecal specimens (dogs 22 and 43) that potentially belong to a *Strongyloides* sp. helminth but could not be confidently assigned to a species given the information available. Two *cox*1 sequences were obtained for dog 22. One of these clustered with a *cox*1 sequences from *Strongyloides mirzai* (GB: AB526307.1), a helminth that infects a Japanese pit viper. The second sequence from dog 22 clustered between the *S. mirzai* clade and the *S. fuelleborni* clade yet also clustered in a position immediately basal to all hookworms (Figure 1). This sequence also obtained a nearest BLASTN hit to a *Necator* sp. sequence (GB: AB793563.1), though its next best hit based on an online BLASTN search was to a sequence from *S. stercoralis* (GB: LC179452.1). Three unique *cox*1 sequences were obtained from dog 43 (GB: MK434220, MK434221; MK434222), one clustering with *S. mirzai* and a second clustering with two sequences of the *Strongyloides* sp. ‘loris’ clade (GB: LC197958.1, LC197946.1). When submitted to an online BLASTN search, the sequence clustering alongside *S. mirzai* also obtained top hits to free living nematodes (*Ektaphelenchus* sp. and *Bursaphelenchus populi*, GB: JX979197.1 and HQ699854.1 respectively), yet several of its top hits were also to *S. fuelleborni cox*1 sequences. The third sequence from dog 43 clustered between the *Strongyloides* sp. ‘loris’ clade and a clade containing all *S. stercoralis* sequences (Figure 1).

### Mixed genotype infections with *Strongyloides stercoralis*

In two samples, dog 6 and dog 18, a complete genotype was obtained (a sequence for *cox*1, HVR-I and HVR-IV), indicating mixed *S. stercoralis* infections. When examining the number of reads that mapped to each haplotype for these specimens (not shown), for dog 6 approximately 20% of reads were assigned to haplotype II and 80% to haplotype IV for the HVR-I region. For the HVR-IV region, approximately 20% of reads were assigned to haplotype A and 80% to haplotype B. With this information it was deduced that this dog was infected with two strains of *S. stercoralis*, one from the human / dog clade (genotype II/A) and another from a dog-specific clade (genotype IV/B). Interestingly, two *cox*1 sequences were obtained from this dog, one assigned to the human / dog clade and another assigned to dog clade 2, supporting our deduction. Dog 18 was also infected with two types of *S. stercoralis*, with a genotype of II + VI / A + E assigned to this specimen. Given that the number of reads assigned to each of these types fell between 40% and 50%, it is difficult to link the HVR-I types identified here to their corresponding HVR IV type. While the specimen from dog 18 possessed two 18S genotypes, evidence was only found for a single *cox*1 sequence. There were two *S. stercoralis* strains found in the HVR-IV region of the dog 13. Approximately 50% of reads were assigned to the haplotype A and 50% to a new haplotype G.

### Detection of non-*Strongyloides* sp. *cox*1 sequences and mixed helminth infections

Twenty-three *cox*1 sequences were attributed to *Ancylostoma* spp. and one dog (dog 6) was infected with an *Ancylostoma* sp. clustering closely with *Anyclostoma ceylanicum* (Figure 1, cyan clade), and two distinct genotypes of *S. stercoralis*. Sequences were obtained from dogs 14, 15 and 16 that belong to a *Metastrongylus*-like helminth, possibly *Metastrongylus salmi*. Two sequences attributed to *Ancylostoma caninum* were also obtained from dog 15, and a fourth sequence belonging to an *Ancylostoma* sp. was also detected in this dog. A sequence was obtained from dog 37 that obtained BLASTN hits to *S. fuelleborni* sequences (e.g., GB: AB526303.1, 86.2% identity). Agglomerative nested clustering placed this sequence in a position basal to all *Strongyloides* and hookworm sequences included in this analysis. This *cox*1 sequence also obtained close BLASTN hits (87% identity) to *Aphelenchoides* sp. (GB: KX356839.1) and *Bursaphelenchus luxuriosae* (AB097863.1) which are free living mycophagous and/or potentially plant parasitic nematodes. The *cox*1 from dog 28 does not appear to be helminth in origin and most closely resembles a *cox*1 sequence from a rotifer; *Macrotrachela quadricornifera* (GB: JX184003.1), which served as a convenient outgroup for the clustering analysis (Figure 1).

### The presence of *Strongyloides* and other helminths in remote Australian communities

*Strongyloides stercoralis* was detected in 10 specimens from seven remote communities in Australia by genotypic analysis. While all specimens included in this study tested positive for *S. stercoralis* using a published real-time PCR assay (Verweij *et al*. 2009), deep sequencing found no evidence of *S. stercoralis* infection in several cases where instead, an infection with another helminth (usually *Ancylostoma* spp.) was confirmed. Some of these sequences clustered closely to *A. ceylanicum* yet appeared to be distinct. However, it should be noted that these sequences were identical to a published sequence assigned to *A. caninum* (GB: AJ407962.1) which did not overlap completely with our amplicon (50 bases short), so we cannot be sure they possess the same sequence type. This necessitates the conservative assignment of these sequences to *Ancylostoma* sp. Most non-*Strongyloides* spp. sequences obtained from dogs were more confidently assigned to *Ancylostoma caninum* (Figure 2). Two dogs from community 2 were also infected with *S. stercoralis* while in one dog from community 5, a cryptic *Strongyloides* sp. possessing a unique *SSU* genotype for both HVR-I and HVR-IV was detected (dog 22, genotype VIII/F). Interestingly, a *Metastrongylus*-like *cox*1 sequence was detected in all dogs from community 3 that were tested, and a single dog (dog 15) from this same community was infected with *A. caninum* and at least one other *Ancylostoma* sp. (Figure 2, cyan clade). We propose that sequences obtained from community 6 are from environmental organisms, possibly representing extraneous contaminants given that one represents a rotifer-like sequence (dog 28) (GB: MK434225) and the other obtained BLASTN hits to free-living nematodes (dog 37) (GB: MK434224). Community 4 and community 7 are represented by a single typed specimen each (dog 18 and dog 45 respectively), that include unique sequences from a helminth that we can confidently assign to the genus *Strongyloides* (Figure 1).

## Discussion and conclusions

In our study we observed that HVR-IV haplotype A is associated with strains infective for both humans and dogs, while HVR-IV haplotype B is restricted to strains that are only infectious to dogs. The same was discovered in a recent study on *S. stercoralis* from humans and dogs in Cambodia, two genetically distinguishable *S. stercoralis* populations were identified based on the HVR-IV region. The HVR-IV haplotype A strain was found to be dog and human infective, while HVR-IV haplotype B strain was shown to be dog specific (Jaleta et al., 2017). Supporting earlier findings, our results also showed that haplotype II of HVR-I is found in combination with haplotype A of the HVR-IV region, and haplotype IV of the HVR I region is only found in combination with haplotype B, which is specific to dogs. One dog possessing the HVR-I haplotype IV (dog 6) had a mixed *S. stercoralis* infection, presumably with worms of the genotype II/A (a type infectious to dogs and humans) and others with the genotype IV/B (a dog specific genotype). The detection of two *cox*1 sequences that cluster in the dog / human and dog specific clades respectively supports this assessment. In an Australian dog, we also identified HVR-I haplotype VI which has only been previously reported in European dogs. Interestingly, this dog (dog 18) also had a mixed genotype infection that included a novel *S. stercoralis* HVR-IV genotype (haplotype E), that was linked to a *cox*1 sequence clustering in the dog / human *S. stercoralis* clade.

In agreement with previous reports, the current study demonstrated that the *SSU* HVR IV region in the *SSU* rDNA can be used to detect within species differences that correspond with the genetic clades that appear when the same specimens are analyzed at the *cox*1 locus (Hasegawa et al., 2016, Jaleta et al., 2017) (Figure 1). To support analysis of the *cox*1 locus by deep sequencing, the *cox*1 PCR assay described here was designed so that merged paired-end Illumina reads span the entire length of the amplicon. This greatly reduces the complexity of *in silico* analysis when mixed *cox*1 genotypes are encountered. A trade-off of using a short amplicon for this analysis is that it may capture less diversity. Additionally, short sequences are of limited use in phylogenetic analysis. However, phylogenies are only truly relevant for constructing the evolutionary history of taxa and because evolutionary analysis was not the objective of this study, agglomerative nested clustering was used to group *cox*1 sequences based on their pairwise sequence identity. Despite its limitations, our *cox*1 assay clearly resolved the dog and human infective *S. stercoralis* genotypes into a clade that is distinct from the dog-specific types, which was our primary objective, and allowed us to compare our results to those obtained in previous studies. Furthermore, we show that the *cox*1 fragment amplified here clearly distinguishes *S. stercoralis*-derived *cox*1 amplicons from other helminth species including *S. fuelleborni* and multiple species of hookworm.

As part of this study, we modified the typing scheme initially reported by Jaleta *et al*. (2017) to include certain *SSU* sequences previously published in GenBank in order to accommodate our novel genotypes. The rationale for these additions was several fold (Table 2 and Table 3). Firstly, we observe that a sequence published in 1993 by Putland *et al*. (1993) (GB: M84229.1) was distinct from other *S. stercoralis* sequences that have been mentioned in the literature since, differing from HVR-IV haplotype A by one SNP at position 125 (Figure 2) (Putland et al., 1993). Consequently, the HVR-IV region of this sequence was added to the typing scheme as haplotype C due to its historic value (Figure 2). The HVR-I sequence of M84229.1 is so drastically different to that of other *S. stercoralis* haplotypes (and to that of any other *Strongyloides* spp. in general), and given that it has not been reported in the literature since its initial publication, it was not added to the typing scheme. Next, we observed that *S. stercoralis SSU* HVR-I haplotype IV (reportedly a dog-specific type) is also found in sequences assigned to *S. procyonis* from a Japanese badger (GB: AB272234.1). To reconcile this observation, we incorporated the HVR-IV region of sequences assigned to *S. procyonis* into the typing scheme, referring to them as haplotype D (Table 3, GB: AB272234.1 and AB205054.1). This also meant that the HVR-I sequence of the *S. procyonis SSU* (GB: AB205054.1) became HVR-I haplotype VII. Hence, the new *SSU* HVR-I genotype from dog 22 (GB: MK468661) became type haplotype VIII. A novel SSU HVR-I genotype from dog 45 was also discovered as part of this study (GB: MK468662), and its sequence was most similar to the *SSU* HVR-I region from *Strongyloides* sp. Okayama, isolated from a Japanese striped snake (GB: LC038066.1). As this sequence was already in GenBank prior to the commencement of this study, the HVR-I region of LC038066.1 was assigned to haplotype IX, while the novel sequence obtained from dog 45 was assigned to haplotype X (GB: MK468662). As haplotypes A to D for HVR-IV had been assigned to other sequences, the novel genotypes discovered in dogs 18, 22 and 13 were assigned to HVR-IV haplotype E, F and G respectively (GB: MK468674, MK468675, and MK468676). Finally, the HVR-IV sequence from *Strongyloides* sp. Okayama (GB:LC038066.1) was assigned to haplotype H because it obtained a nearest match to the HVR-IV regions sequenced from dog 45, which was consequently assigned to haplotype I (GB: MK468677). Also note that all HVR-I and HVR-IV types discussed above (both novel and previously published) are more similar to each other than they are to the corresponding *SSU* regions from *S. ratti*. Consequently, they do not provide enough information on their own to make confident species assignments (Figure 2).

Ultimately, while the typing scheme developed by Jaleta *et al*. (2017) was originally designed to consider *S. stercoralis* genotypes alone, the detection of several novel cryptic genotypes that: (1) cannot be confidently assigned to a species, (2) are nonetheless greater than 99% similar to each other and to known *S. stercoralis* genotypes and (3), are genotypes detected in the same host (dogs), means that the adjustments made here represent the most straightforward solution to the issue at hand. Moreover, being an isolated continent, it is possible that Australian dogs might be infected with genetically distinct *S. stercoralis* strains (Cawood and Korsch, 2008). Consequently, it is not unreasonable to suggest that some of the cryptic dog *Strongyloides* genotypes described herein (i.e., from dogs 18, 22, 13 and 45) might be attributable to truly novel *S. stercoralis* genotypes restricted to Australia. However, a larger sample number is needed along with additional sequences and morphological analysis of multiple specimens before these sequences can be assigned to a species of helminth. Similarly, it is plausible that the *cox*1 sequences obtained from dog 43 are attributable to a helminth belonging to the genus *Strongyloides*, yet the lack of any *SSU* sequences associated with this specimen makes it difficult to draw any solid conclusions in that regard.

This study employed an alteration of the Jaleta *et al*. (2017) genotyping assay developed at the Centers for Disease Control and Prevention (J. Barratt) for adaptation to NGS technologies. The assay was designed to genotype *Strongyloides* sp. and potentially detect mixed helminth infections (e.g., hookworm and *Strongyloides*) when applied directly to DNA extracted from faeces and other biological specimens. This method has great advantages over previous genotyping techniques in that it can be undertaken directly from faecal DNA extracts and does not require culture of larvae for extraction of DNA from individual worms. Furthermore, the depth of sequencing provided by NGS allows the detection of all genotypes in a single sample (Zahedi et al., 2017).

This study had a number of limitations including the collection of dog stool from the environment where they could possibly have been contaminated with extraneous environmental organisms or their DNA. As noted in Table 1, the *cox*1 PCR employed in this study also detects multiple hookworm species, and was even found to amplify the *cox*1 sequence of a *Metastrongylus*-like helminth. This may represent an advantage of the method if simultaneous detection and genotyping of multiple pathogenic intestinal nematodes from dogs and humans is required. However these results should be viewed with caution. Given the sensitivity of deep sequencing, we suspect that detection of *cox*1 sequences resembling those of *Metastrongylus salmi* could be attributable to the consumption of pig offal by dogs in community 3. While *Metastrongylus* sp. are known to occasionally infect other species including humans (Calvopina et al., 2016), the genus is generally thought to be specific to pigs. We also note that the genotype from dog 45 closely resembles that of a reptile-infecting *Strongyloides* sp. and its presence in a dog could be due to consumption of reptiles or reptile feces by the dog. Consequently, these cases may represent an incidental finding rather than true infections with a *Metastrongylus*-like helminth or a *Strongyloides* sp. resembling those found in reptiles. Similarly, the *cox*1 assay described here detected DNA from potentially free – living nematodes. A sequence obtained from dog 37 received a BLASTN hit to *S. fuelleborni* (GB: AB526306.1), though with only 86.2% identity. However, agglomerative nested clustering placed this sequence in a position basal to all *Strongyloides* and hookworm sequences included in this analysis. This sequence also obtained close BLASTN hits (87% identity) to *Aphelenchoides* sp. (GB: KX356839.1) and *Bursaphelenchus luxuriosae* (AB097863.1) which are mycophagous and/or potentially plant parasitic nematodes. This sequence could therefore represent a free-living nematode that came into contact with the fecal specimens in the environment between when the stool was passed and collected. Surprisingly, a sequence similar to one obtained from a rotifer; a free-living extremely distant relative of nematodes, was detected in the specimen from dog 28 using this assay. This also likely represents contamination of the stool specimen from the local environment prior to its collection.

Several unique sequences assigned to *Strongyloides* spp., *Ancylostoma* spp., *Metastrongylus* sp. and some unknown helminth species were detected for the first time using the technology described here. We propose that some of the cryptic *Strongyloides* sp. genotypes we encountered are potentially unique to the Australian continent and may have diverged from southeast Asian *Strongyloides* populations as a result of vicariance. Discovery of a *cox*1 sequence that clusters most closely to a *Strongyloides* sp. identified from a slow loris might support this (dog 43, GB: MK434221), given that lorises are endemic to southeast Asia. Aside from these cryptic types, several examples of the classic *S. stercoralis* genotypes described by Jaleta *et al*. (2017) were detected. Consequently, our study was able to independently support previous reports of at least two genetically distinct groups of *S. stercoralis*; one infecting both dogs and humans and another group that is specific to dogs. While this does not prove the potential of dogs as a reservoir of *S. stercoralis*, it supports the hypothesis of zoonotic transmission in remote Australian communities. As discussed previously and with respect to the One Health approach (Rock et al., 2009), we suggest that humans and dogs should be treated concomitantly in these communities to control strongyloidiasis (Beknazarova et al., 2017b, Jaleta et al., 2017). Ultimately, we confirm for the first time that potentially zoonotic *S. stercoralis* genotypes are present in Australia and suggest that dogs represent a potential reservoir of human strongyloidiasis in remote Australian communities.

## Author Contributions

MB: Designed study, obtained ethics approvals, collected specimens, extracted DNA, ran real-time PCR, optimized and ran PCR assays for *SSU* HVR-I, HVR-IV and *cox*1, performed Illumina library preparation, performed *in-silico* analysis, data analysis and interpretation, wrote and reviewed draft manuscripts; JB: Designed PCR assay for *SSU* HVR-I, HVR-IV and *cox*1, performed *in silico* analysis and data interpretation, constructed cluster dendrogram, wrote and reviewed manuscript drafts; RB: Designed study, reviewed manuscript drafts, obtained funding, reviewed manuscript drafts; ML: Carried out PCR product normalization, prepared Illumina libraries for deep sequencing, and sequenced amplicons; HW: Provided PhD supervision of MB, designed study, obtained ethics approvals, obtained funding, reviewed manuscript drafts; KR: Provided PhD supervision of MB, designed study, obtained ethics approvals, obtained funding, reviewed manuscript drafts.

## Acknowledgments

The authors would like to thank Professor Robert W. Baird and Dr Richard Sullivan at the Royal Darwin Hospital, Darwin Pathology, NT and Dr Gemma Robertson at the Health Support Queensland, QLD for providing us human *S. stercoralis* DNA samples We would also like to acknowledge the contribution of Animal Management in Rural and Remote Indigenous Communities, Environmental Health Branch at the Department of Health, NT and all the other lovely veterinarians across Australia for helping us collecting dog faecal samples in the remote communities. We acknowledge the help of Dr Rogan Lee, Dr Matthew Watts, John Clancy and Vishal Ahuja at the Westmead Hospital, NSW with sending us *S. ratti* infected rat faeces. We wish to thank staff at the Centers for Disease Control and Prevention in Atlanta, USA. The work has been supported by the Australian Government Research Training Program Scholarship and Flinders University Overseas Travelling Fellowship.

## Supporting Information Legends

**S1 Table. GenBank accession numbers of the SSU HVR I and IV and *cox*1 haplotypes**. The GenBank accession numbers are provided for the SSU HVR I and IV haplotypes generated in this study (MK468654-MK468677). The GenBank accession numbers are provided for the cox1 haplotypes generated in this study (MK434217- MK434258).

